# Western diet-induced shifts in the maternal microbiome are associated with altered microRNA expression in baboon placenta and fetal liver

**DOI:** 10.1101/2022.05.18.492490

**Authors:** Kameron Y. Sugino, Ashok Mandala, Rachel C. Janssen, Sunam Gurung, MaJoi Trammell, Michael W. Day, Richard S. Brush, James F. Papin, David W. Dyer, Martin-Paul Agbaga, Jacob E. Friedman, Marisol Castillo-Castrejon, Karen R. Jonscher, Dean A. Myers

## Abstract

Maternal consumption of a high-fat, Western-style diet (WD) disrupts the maternal/infant microbiome and contributes to developmental programming of the immune system and nonalcoholic fatty liver disease (NAFLD) in the offspring. Epigenetic changes, including non-coding miRNAs in the fetus and/or placenta may also underlie this risk. We previously showed that obese nonhuman primates (NHP) fed a WD during pregnancy results in the loss of beneficial maternal gut microbes and dysregulation of cellular metabolism and mitochondrial dysfunction in the fetal liver, leading to a perturbed postnatal immune response with accelerated NAFLD in juvenile offspring. Here, we investigated associations between WD-induced maternal metabolic and microbiome changes, in the absence of obesity, and miRNA and gene expression changes in the placenta and fetal liver. After ∼8-11 months of maternal WD feeding (mWD), dams were similar in body weight but exhibited mild, systemic inflammation (elevated CRP and neutrophil count) and dyslipidemia (increased triglycerides and cholesterol) compared with dams fed a control diet. The maternal gut microbiome was mainly comprised of *Lactobacillales* and *Clostridiales*, with significantly decreased alpha diversity (*P* = 0.0163) in WD-fed dams but no community-wide differences (*P* = 0.26). At 0.9 gestation, mRNA expression of *IL6* and *TNF* in mWD-exposed placentas trended higher, while increased triglycerides, expression of pro-inflammatory *CCR2*, and histological evidence for fibrosis were found in mWD-exposed fetal livers. In the mWD-exposed fetus, hepatic expression levels of miR-204-5p and miR-145-3p were significantly downregulated, whereas in mWD-exposed placentas, miR-182-5p and miR-183-5p were significantly decreased. Notably, miR-1285-3p expression in the liver and miR-183-5p in the placenta were significantly associated with inflammation and lipid synthesis pathway genes, respectively. *Blautia* and *Ruminococcus* were significantly associated with miR-122-5p in liver, while *Coriobacteriacea* and *Prevotellacea* were strongly associated with miR-1285-3p in the placenta; both miRNAs are implicated in pathways mediating postnatal growth and obesity. Our findings demonstrate that mWD shifts the maternal microbiome, lipid metabolism, and inflammation prior to obesity and are associated with epigenetic changes in the placenta and fetal liver. These changes may underlie inflammation, oxidative stress, and fibrosis patterns that drive NAFLD and metabolic disease risk in the next generation.

## Introduction

Nonalcoholic fatty liver disease (NAFLD) is the most common liver disease worldwide. Characterized by simple steatosis (excess liver fat), NAFLD may progress to nonalcoholic steatohepatitis (NASH) with inflammation and fibrosis, leading to cirrhosis and increased risk for hepatocellular carcinoma (1). The CDC estimates that >18 million women of reproductive age in the U.S. are obese, and maternal obesity is strongly linked to inflammatory and metabolic disorders in the offspring (2-7). Alarmingly, 1 in 5 preschoolers are obese (8) and 1/3 of obese youth are diagnosed with NAFLD (9). Despite evidence that maternal overnutrition adversely influences metabolic health in human (10) and animal (11-15) offspring, there is a fundamental lack of insight into molecular mechanisms by which maternal dietary exposures reprograms fetal immune development and NAFLD in utero, particularly in models that reflect the human condition.

The placenta acts as the primary interface between mother and fetus, allowing nutrient and oxygen transfer which supports fetal growth and development. Obesity-associated maternal-fetal inflammation contribute to various adverse pregnancy outcomes including placental dysfunction (16, 17), preeclampsia, neurodevelopment (18), intrauterine growth restriction (19, 20), and preterm labor (21, 22). An investigation of the chronic pro-inflammatory milieu in placentas from obese pregnancies showed a two-to three-fold increase in resident macrophages (CD68+ and CD14+) and expression of pro-inflammatory cytokines compared with placentas from lean pregnancies (23). However, the underlying causes of placental inflammation remain unclear. One possible mechanism is through inflammatory metabolites, such as lipopolysaccharides, which are produced by the microbiome, reach higher systemic levels in obesity (24), and can directly interact with placental Toll-like receptor 4 (25). The role of a Western-style diet (WD) versus maternal obesity in disruption of placental function and inflammatory state remains to be elucidated, as most animal models of maternal obesity have relied upon a WD to attain and maintain obesity.

The maternal gut microbiome changes during pregnancy and is influenced by maternal diet, maternal obesity, excessive gestational weight gain, and gestational diabetes mellitus (GDM) (26-28). Animal and human studies (29-32) support a role for the maternal gut microbiome and bacterial metabolites in the pathophysiological changes accompanying maternal obesity (33), offspring NAFLD (34, 35), and infant inflammatory disorders (36). WD-induced maternal obesity is associated with impaired microbiome and placental structural and functional changes, concomitant with oxidative stress and inflammation in the intrauterine environment (37-39). Trophoblasts recognize and respond to bacterial products (40) and integrate microbial-derived signals via pathways mediating the response to Toll-like receptors or epigenetic modifications (41). Although mechanisms by which products of gut dysbiosis affect placental inflammation/function and fetal inflammation are not well understood, reduction in beneficial bacterial metabolites might play a role.

Emerging studies are addressing the association of maternal diet and obesity with alterations to epigenetic signatures and microbiome function in offspring (13, 42). The epigenome consists of DNA and histone modifications that produce heritable changes in transcription and cellular function (43). One form of epigenetic modification are microRNAs (miRNAs). MiRNAs are small, noncoding RNAs that bind to the 3’ untranslated region of protein-coding mRNAs, degrading or repressing translation of the targeted mRNAs. In humans, miR-122, miR-34a, miR-21, and miR-29a were shown to regulate lipid metabolism, oxidative stress, and inflammation in the liver, with a crucial role in the pathophysiology of NAFLD (44-47). Studies investigating the impact of early changes in miRNAs in the fetal liver are sparse; however, using large-scale sequencing and pathway analysis in baboon fetal liver, Puppala et al. identified 11 miRNAs targeting 13 genes in metabolic pathways (TCA cycle, oxidative phosphorylation, and glycolysis), the proteasome and WNT/β-catenin signaling (48). In the placenta, miRNA expression levels are disrupted by hypoxia (49), maternal exposure to toxic agents such as cigarette smoke (50) and bisphenol A (51), and GDM (52). In humans, elevated pre-pregnancy BMI has been associated with lower expression levels of placental miRNAs; these differences were modified by offspring sex and maternal gestational weight gain (53). However, studies investigating associations between maternal diet, SCFAs, and epigenetic modifications in the placenta and fetal liver are lacking. Here, we leveraged our well-established Olive baboon (*Papio anubis*) nonhuman primate (NHP) model of maternal WD consumption to investigate changes in the maternal microbiome and their association with placental and fetal liver miRNA expression profiles in non-obese pregnancy.

## Materials and Methods

### Animal model

All experiments utilizing baboons were performed in compliance with guidelines established by the Animal Welfare Act for housing and care of laboratory animals as well as the U.S. National Institutes of Health Office of Laboratory Animal Welfare Public Health Service Policy on Humane Care and Use of Laboratory Animals. All animals received environmental enrichment. All experiments were conducted in accordance with and with approval from the University of Oklahoma Health Sciences Center Institutional Animal Care and Use Committee (IACUC; protocol 302043 #22-025-AH).

Nulliparous Olive baboon females (*Papio anubis*; n=11; 4-7 years old; puberty is ∼4-5 years of age) with similar lean body scores were randomly separated into two primary cohorts with similar mean age and body weight, taking into consideration social stratification. Dams were housed in diet groups (n=5/6 per corral) for the duration of the study. Control diet (CD) dams were fed standard monkey chow diet (5045, Purina LabDiets, St. Louis, MO; 30.3% calories from protein, 13.2% from fat, 56.5% from carbohydrates) and the WD dams were fed a high-fat, high-simple sugar diet (TAD Primate diet, 5L0P, Purina; 18.3% calories from protein, 36.3% of calories from fat, 45.4% from carbohydrates), matched to CD for micronutrients and vitamins, and supplemented with continuous access to a high-fructose beverage (100 g/L KoolAid™). Both groups were provided the same daily enrichment foods (fruits and peanuts). The TAD diet/high-fructose drink is widely used to study the role of an excess energy intake, high-saturated fat, high-fructose diet on physiological systems in NHPs (54, 55) and is consistent with human WDs. After an initial 3 months of WD to allow for collection of baseline samples and acclimation to WD, dams were bred to males that had been previously fed CD. Blood, fecal samples, and anthropometric measurements (body weight and sum of skin folds) were obtained under ketamine (10-20 mg/kg) and acepromazine (0.05-0.5 mg/kg) sedation administered via intramuscular injection. Following chemical sedation, cephalic or saphenous vein catheters were placed for blood draw. At 0.6 gestation (G; term is ∼183 days gestation) baboons were fasted overnight. Intravenous glucose tolerance test (IVGTT) was performed under ketamine and acepromazine sedation and two appropriately sized intravenous catheters were placed into each saphenous or cephalic vein, or a combination thereof, one for infusion of dextrose and one for venous blood collection. A baseline blood draw was taken from one catheter and an intravenous bolus of 50% dextrose (0.5 g/kg body weight) was administered over 30 seconds through the second catheter. Blood glucose was measured in venous blood using a glucometer at time 0, 2, 4, 8, 12, 16, 20, and 40 min post dextrose infusion.

### Maternal blood analyses

Complete blood counts (CBCs) were obtained for the dams from EDTA-anticoagulated whole blood samples. CBCs included analyses for red blood cells, white blood cell count (neutrophils, lymphocytes, monocytes, eosinophils and basophils), platelets, and hemoglobin. Maternal serum samples collected at 0.6 G were analyzed for C-reactive protein (CRP) using an hsCRP ELISA kit (MP Biomedicals, Solon, OH) according to the manufacturer’s protocol with 1:100 serum dilution and were analyzed for IL-6 using an old world monkey IL-6 ELISA kit (U-CyTech Biosciences, the Netherlands) according to the manufacturer’s protocol. Maternal serum samples were analyzed for triglycerides (TGs) using a triglyceride colorimetric assay kit (Cayman Chemical, Ann Arbor, MI) according to the manufacturer’s protocol. High-density lipoprotein (HDL) and low-density lipoprotein/very low-density lipoprotein (LDL/VLDL) levels were quantified in serum samples taken at 0.6 G from fasted dams using EnzyChrom HDL and LDL/VLDL Assay Kits (BioAssay Systems, Hayward, CA). Samples were assayed in duplicate according to the manufacturer’s instructions.

### Cesarean section

At 0.9 G, dams were anesthetized using isoflurane and fetuses were delivered by cesarean section. Fetal and placental weights were obtained, and placental and liver tissue samples were processed for histology or flash-frozen in liquid nitrogen and stored at -80° C for subsequent analyses.

### Liver histology

Fetal liver tissue samples from the left lobe were fixed in 10% formalin for 24 h followed by storage in 70% EtOH. Histology was performed by the OUHSC Stephenson Cancer Tissue Pathology Core. In brief, samples were paraffin-embedded and sectioned for H&E and picrosirius red staining. Fresh-frozen liver from the left lobe was fixed in OCT compound, sectioned, and fixed with formalin for 5 min and washed with PBS. LipidSpot lipid droplet stain (Biotium, Fremont, CA) was added to the sections and incubated for 20 min. Sections were washed with PBS, counterstained with DAPI, and mounted using VectaMount AQ aqueous mounting medium (Vector Labs, Burlingame, CA). Slides were visualized using a Cytation 5 microscope and Gen5 imaging software (Agilent, Santa Clara, CA).

### Placenta immunofluorescence

Immediately upon delivery of the placenta during cesarean section, placental tissue was dissected from each cotyledon: one half of each sample was flash-frozen and stored at -80°C and the other half was fixed in 4% paraformaldehyde for 48 h, transferred to 70% EtOH, and paraffin embedded. Thin sections (5-micron thickness) were obtained every 150 microns from the paraffin blocks and placed onto slides. For immunofluorescence (IF) labelling, sections were selected to allow for a total of four sections per cotyledon at a minimum of 150 microns between sections. Slides were baked for 1 h at 56°C, deparaffinized and antigen retrieval was performed in a Retriever 2100 instrument with R-Universal epitope recovery buffer (Electron Microscopy Sciences, Hatfield, PA). After retrieval, slides were blocked in 5% normal donkey serum for 1 h, then primary antibody (MAC387 [anti-S100A9 + Calprotectin], Abcam, Cambridge, UK) was added and slides were incubated for 16 h at 4°C with humidification. Slides were subsequently allowed to equilibrate to RT for 1 h. Secondary antibody (donkey anti-mouse IgG F(ab’)2 Alexa Fluor 594, Jackson ImmunoResearch, West Grove, PA) was added and slides were incubated for 1 h, covered, at RT. Slides were counterstained for 5 min with DAPI and coverslipped using Shur/Mount. Slides were visualized using a fluorescence microscope (Olympus BX43) and images were captured using CellSens imaging software (Olympus). Four 150×150 micron images were selected randomly per tissue section (700×500 microns) with no overlap between selected sections and macrophages were counted by a researcher blinded to the treatment group. Total macrophages were summed for all images per placenta and scored.

### Fetal liver and placental tissue analyses

RNA was extracted from flash-frozen fetal liver (left lobe) samples using a Direct-zol RNA miniprep kit (Zymo Research, Irvine, CA)) per manufacturer instructions. cDNA was synthesized from 1 ug RNA using iScript Supermix (Bio-Rad, Hercules, CA) per manufacturer instructions. Gene expression was measured using real time qPCR with PowerUp SYBR Green Master mix on a QuantStudio 6 instrument (Thermo Fisher Scientific, Waltham, MA). Results were normalized to *RPS9* (ribosomal protein S9) using the comparative Ct method. For placenta, qPCR was performed similar to fetal liver except using a CFX96 RT-PCR Detection System (Bio-Rad, Hercules, CA). Placental results were normalized to *ACTB* using the comparative Ct method. Primers for qPCR are shown in **Supplementary Table S1**. Fetal liver TGs were extracted as described previously (32) and quantified using Infinity Triglycerides Reagent (Thermo Fisher) with normalization to starting tissue weight.

Total RNA, including miRNA, was isolated from flash-frozen fetal liver (left lobe) and placenta tissues (two samples/placenta from separate cotyledons) using miRNeasy mini kit (Qiagen, Germantown, MD) following manufacturer’s instructions. TaqMan MicroRNA Assays (Thermo Fisher) were used for reverse transcription and real-time qPCR (miR-122-5p, assay ID 002245; miR-204-5p, assay ID 000508; miR-34a-5p, assay ID 000426; miR-21-5p, assay ID 000397; miR-183-5p, assay ID 000484; miR-29a-3p, assay ID 002112; miR-185-5p, assay ID 002271; miR-145-3p, assay ID 002149; miR-1285-3p, assay ID 002822; miR-199a-5p, assay ID 000498; miR-182-5p, assay ID 000597). Reverse transcription was carried out using TaqMan MicroRNA Reverse Transcription kit (Thermo Fisher), an RT primer from a specific TaqMan MicroRNA Assay, and 10 ng total RNA following manufacturer’s instructions. The miRNA-specific cDNA templates were placed on ice and used immediately for qPCR or stored at -20°C for 1-2 days prior to use. Real-time PCR reactions were performed in duplicate using TaqMan Universal Master Mix II, no UNG (Thermo Fisher) and the corresponding TaqMan MicroRNA Assay following manufacturer’s instructions. Assays were performed on a QuantStudio 6 Real-time PCR System. Ct values were calculated and the relative miRNA expression levels were quantitated with the comparative Ct method and using miR-92a-3p (assay ID 000431) for normalization.

### SCFA analysis

Frozen feces (100-200 mg) were added to a vial continuing 200 μg/L of deuterated butyric acid (internal standard), 0.2 g/ml NaH_2_PO_4_, and 0.8 g/ml ammonium sulfate (adjusted to pH 2.5 with phosphoric acid). External standards were prepared with 0.2 g/ml NaH_2_PO_4_, 0.8 g/ml ammonium sulfate, 200 μg/L deuterated butyric acid (internal standard), 200 μg/L 2:0 (acetic acid), 100 μg/L 3:0, and 50 μg/L of 4:0, 5:0, and 6:0 (adjusted to pH 2.5). All vials were quickly capped and vigorously agitated. A 7890A gas chromatograph equipped with 7697A headspace sampler, 5975C mass spectrometer detector, and DB-FATWAX UI 30 × 0.25 × 250 column was used for analysis (Agilent). The headspace sampler oven, loop, and transfer line temperatures were held at 50°C, 100°C, and 110°C, respectively. The vial equilibration time was 30 min and the injection duration was 1 min. The vial fill pressure was 15 psi and the loop final pressure was 1.5 psi. Loop equilibration time was 0.05 min. The GC inlet and MS interface were held at 250°C. The oven temperature was held at 120°C for 2 min, ramped at 5°C /min to 140°C, ramped at 20°C/min to 220°C, and held at 220°C for 1 min. Helium carrier gas flowed constantly at 1.2 ml/min and the split ratio was 10:1. SCFAs were detected in SIM ion mode using *m/z* values of 43, 45, 60, 63, 73, 74, 77, and 87. Quantities of 2:0, 3:0, 4:0, 5:0, and 6:0 were determined by comparison to external standards.

### Microbial DNA extraction and sequencing

DNA extraction from feces from dams employed the DNeasy PowerSoil Pro Kit (Qiagen) per manufacturer’s instructions with the following modification. Aliquots of 300 mg of fecal material were weighed, resuspended in 800 μL Solution CD1, and incubated at 60°C for 10 min. Library construction and DNA sequencing were performed by the OUHSC Laboratory for Molecular Biology and Cytometry Research. Library construction employed the Nextera XT Library Preparation Kit (Illumina, San Diego, CA). Samples were barcoded for multiplexing, and sequenced on an Illumina MiSeq using paired-end sequencing with a 600 cycle MiSeq Reagent Kit v3 (Illumina).

### Data Analysis

Maternal data, fetal liver TGs, and qPCR data were analyzed for comparisons between CD and WD using an unpaired, 2-tailed Student’s *t* test; significance was determined as *P* < 0.05. Initial processing of raw microbiome data employed QIIME2 2019.10 software (56). Sequenced 16S rRNA raw fastq files were imported as demultiplexed paired end reads with a Phred score of 33. Sequences were trimmed, quality filtered and denoised into amplicon sequence variants (ASVs) using DADA2 (57). ASVs were then aligned de novo using MAFFT (58) and structured into a rooted phylogenetic tree using FastTree2 (59). Alpha diversity (e.g., Faith’s Phylogenic Diversity, Shannon) and beta diversity (e.g., Weighted UniFrac distance, Bray-Curtis) were compared between diet groups using Kruskal-Wallis and PERMANOVA, respectively. Taxonomy was assigned to each ASV using a sklearn-based Naïve Bayes taxonomy classifier (60) pre-trained on the Greengenes 13_8 99% OTUs reference database sequences (61). Linear discriminant analysis effect size, LEfSe (62), assessed the raw taxonomic abundance table for significant taxa which are differentially abundant in the context of the experimental groups.

#### Variable selection methods

Due to the small sample size (*n* = 11) and relatively large number of comparison variables for mother (11 measures) and fetus (63 measures), we performed Least Absolute Shrinkage and Selection Operator (LASSO) regularization (63) to utilize the technique’s variable selection properties before testing for significance between maternal and fetal measurements. Briefly, LASSO regression applies a shrinkage term (lambda) to coefficients in the model in order to improve (i.e., reduce) the model’s mean squared error. This type of penalty can reduce the value of some coefficients to zero, eliminating them from the model and providing a method of variable selection. To implement this procedure, we used the R package glmnet (64). Maternal and fetal measures were standardized around the mean and standard deviation before building each model. Each fetal measure was used as the response variable and maternal measures as the explanatory variables; we included an interaction term between the maternal measures and fetal sex for each maternal measure within the model. Since complete data are needed for LASSO regularization, each model was estimated using only samples with complete data (minimum *n* = 5, median *n* = 7) and the best lambda value was selected based on the model with the lowest mean squared error. Maternal variables selected by the LASSO procedure were then run in univariate Anova models against the fetal variable using the full dataset (*n* = 11), *P* values were compiled and FDR correction was applied using the Benjamini-Hochberg procedure. A similar method was applied to compare the fetal measures to each other, however, since only 4 fetal samples had complete data, the fetal measures were broken into three separate datasets: miRNA (33 measures, both liver and placenta), fetal mRNA levels (27 measures), and other measures (fetal weight, heart weight, and liver TGs); fetal sex was included as an interaction term for all models. Comparisons between the fetal miRNA were performed using the liver miRNA measures as the response variable and placental miRNA as the predictor. After breaking up the fetal dataset, we performed the following comparisons: fetal miRNA vs. mRNA, miRNA vs. other measures, and mRNA vs. other measures; sample sizes used for each LASSO procedure reached a minimum of 5, median of 7 datapoints for these comparisons.

LASSO regularization was also used for variable selection to compare the maternal microbiota (*n* = 10) to maternal and fetal measures. To compare the maternal measures to the maternal microbiome, we utilized the R package mpath (65) to perform the LASSO procedure using a negative binomial regression model. Briefly, the microbiota were classified to the taxonomic family and genus levels and filtered to include taxa which were present in at least 50% of the samples and reached at least 1% relative abundance. The microbiota count data were used for input into the glmregNB function for LASSO regularization with default values, and the best model was selected using the Bayesian information criterion. Again, only complete sets of data were used in the LASSO procedure (*n* = 6). Variables selected for each taxon were run in a univariate negative binomial regression with the R package MASS (66) using the full dataset (*n* = 10). *P* values were compiled and FDR correction was applied using the Benjamini-Hochberg procedure. LASSO models were constructed using only complete datasets, while statistics were run on the full dataset. To compare fetal measures as an outcome of the maternal microbiota abundances, we used arcsin square root transformation of the microbiota relative abundance data (67) as predictors for the infant measures. LASSO regularization was used to build these models using glmnet and univariate Anova models were used to test for significance as described in the previous paragraph. The samples sizes used for LASSO regularization between maternal microbiota and infant measures are as follows: fetal miRNA, *n* = 8; fetal mRNA, *n* = 6; and other fetal measures (weight, heart weight, and liver TGs), *n* = 8. Fetal sex was included as an interaction term within these models.

## Results

### WD induces inflammation in dams, placenta, and fetuses and impairs lipid metabolism

The time (days) from initiation of WD to IVGTT (carried out at 0.6 G) ranged from 276 to 480 days (332 ± 39 d); the time from initiation of WD to cesarean section (carried out at 0.9 G) ranged from 324 to 535 days (386.5 ± 39 d). Despite this duration of exposure to WD/high fructose, at 0.6 G, no significant difference in maternal body weight (**Fig. 1A**) or in adiposity index (sum of skin fold thickness, **Fig. 1B**) was observed. These findings are consistent with those reported by the Nathanielsz lab (68, 69) using a similar diet and baboon model, demonstrating that attainment of maternal obesity requires a minimum of 9 mo. to ∼3 years of WD feeding. Although WD-fed (mWD) dams did not become obese, serum CRP (**Fig. 1C**) and neutrophils (**Fig. 1D**) increased, without a change in serum IL-6 (**Fig. 1E**), indicative of mild, systemic inflammation in response to WD, prior to attaining obesity. WD-fed dams also exhibited elevated serum TGs (**Fig. 1F**). Serum HDL cholesterol and LDL/VLDL cholesterol were significantly higher in mWD dams compared with CD-fed (mCD) dams; however, no significant change in total cholesterol was observed (**Fig. 1G**). Blood glucose tolerance tests did not show a significant effect of diet (**Fig. 1H**). Together, these findings demonstrate that a relatively short exposure to WD during pregnancy induces significant systemic inflammation and impaired lipid metabolism in baboon dams prior to a significant change in adiposity or insulin sensitivity.

**Figure 1.**
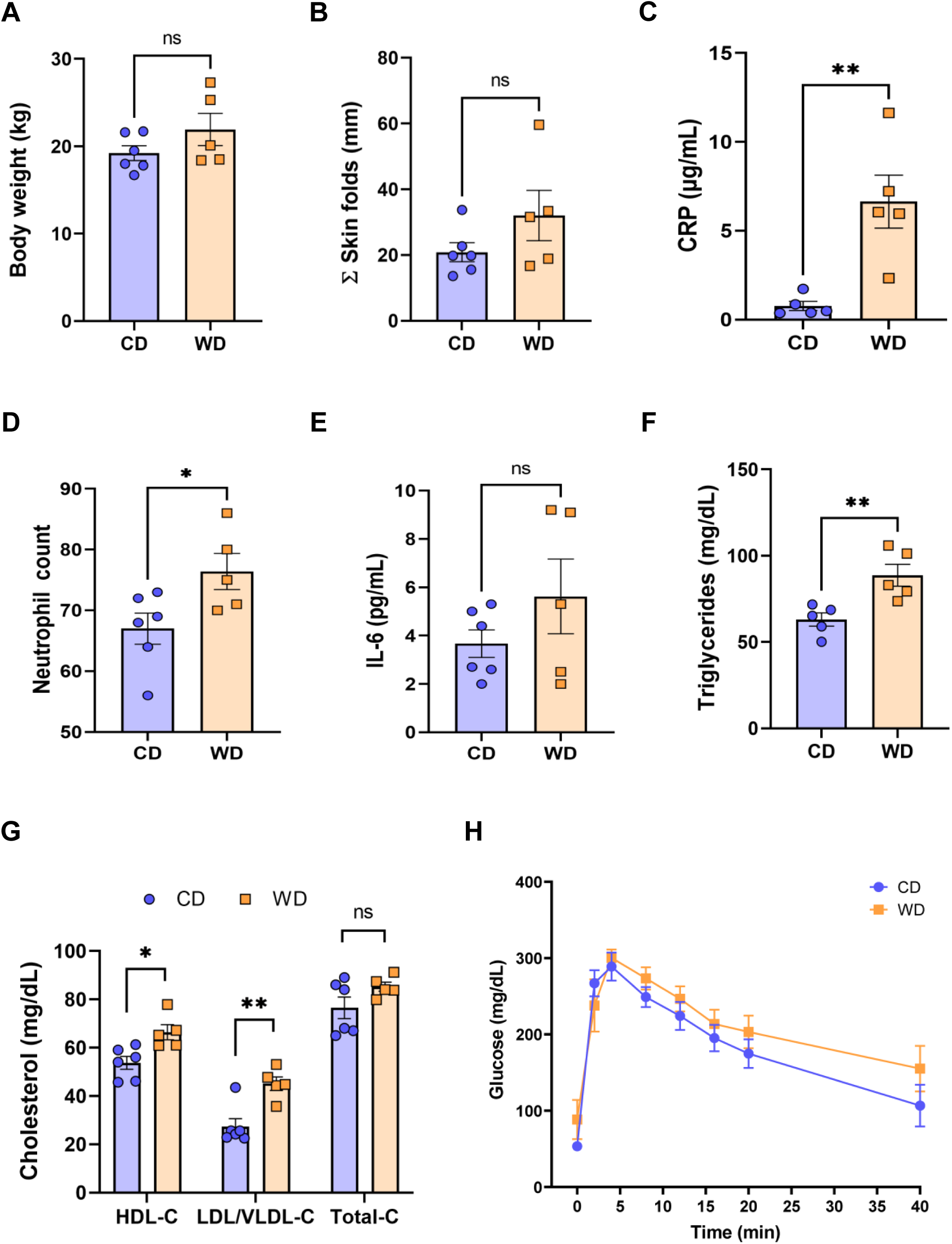
WD-fed dams exhibit alterations in inflammation and lipid metabolism at 0.6 gestation. Maternal body weight (**A**), sum of (Σ) skin folds as a measure of adiposity (**B**). Maternal serum levels of C-reactive protein (CRP, **C**), neutrophil count from complete blood count (**D**), serum IL-6 levels (**E**), and triglycerides (**F**). Cholesterol analysis of red blood cells (**G**) and IVGTT analysis (**H**). *n* = 5-6 CD and *n* = 5 WD. Unpaired 2-tailed Student’s *t* test was used to test significance. \**P* < 0.05, \*\**P* < 0.01.

Given the markers for inflammation observed in dams on WD, we performed immunofluorescence on fixed placental sections obtained at cesarean section using an antibody to MAC387 to label infiltrating monocytes/ macrophages. We observed very few MAC387+ macrophages in mCD placentas (**Fig. 2A**). Macrophage infiltration was more variable in the placenta of mWD dams, with notable macrophages evident in 3 of 5 placentas whilst macrophage presence in two placentas was comparable with those of mCD placentas (**Fig. 2A, right panel**). Expression of cytokines/chemokines (*IL1B, IL6, TNF, IL8*) in placental tissue (**Fig. 2B**) were elevated in mWD placenta but differences did not reach significance when compared with mCD placenta. However, both *IL6* (*P* = 0.08) and *TNF* (*P* = 0.068) exhibited trends for significance. No notable gross placental pathologies (calcifications, infarcts) were observed in mWD placentas compared with mCD placentas.

**Figure 2.**
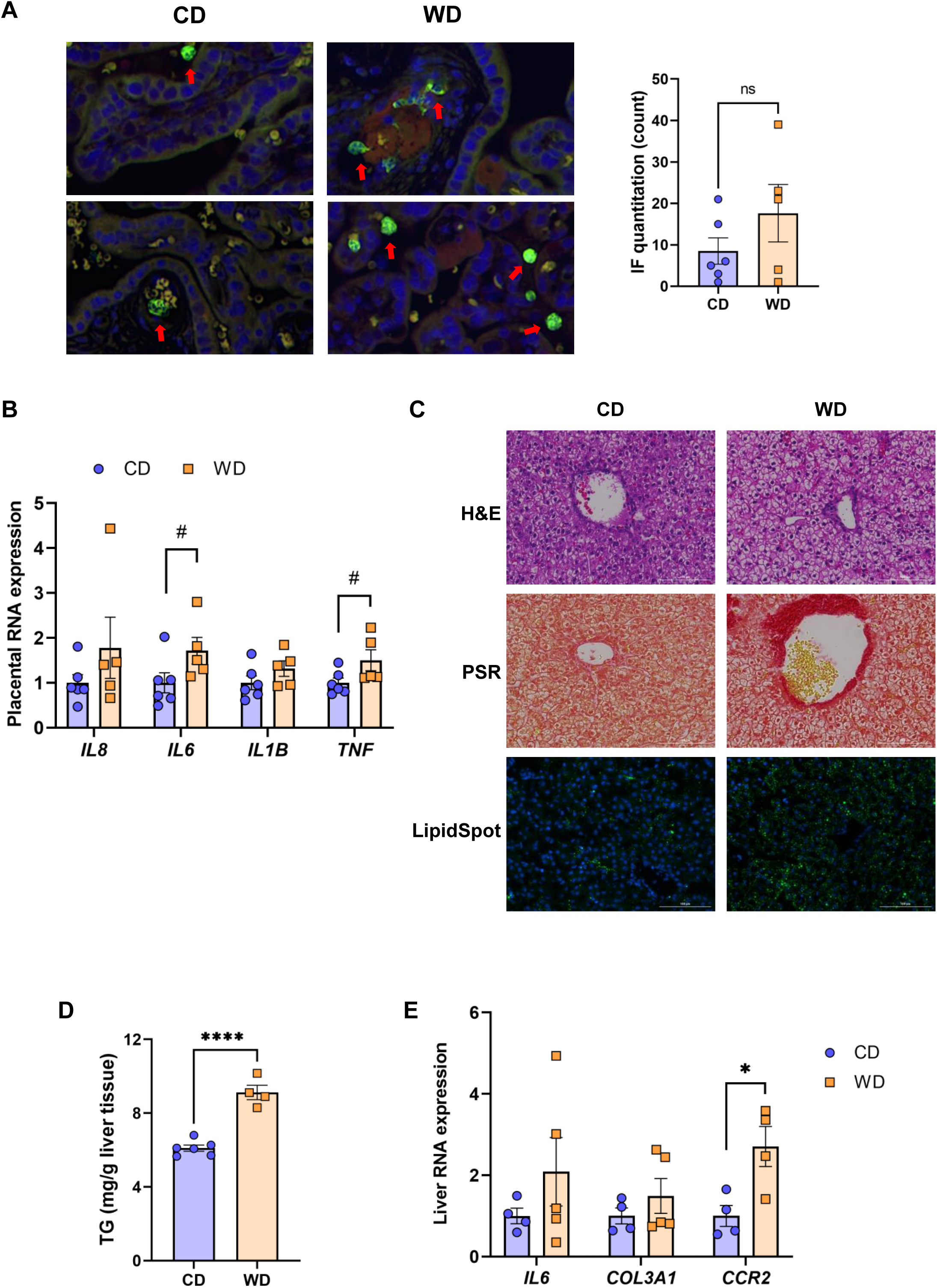
WD exposure increases monocyte infiltration of the placenta and induces fetal hepatic steatosis and fibrosis. Representative images for immunofluorescence in placenta tissue and quantitation (**A**). Red arrows point to MAC387-labeled macrophages (green). Blue staining - DAPI. (**B**) mRNA expression of cytokines in placenta using qPCR. *ACTB* was used for reference. Representative images of histological analysis of fetal liver tissue (**C**) with H&E, picrosirius red (PSR), and LipidSpot, taken at 100 um. Fetal liver triglycerides (TG) (**D**) and mRNA expression analysis using qPCR (**D**) with *RPS9* used for normalization. *n* = 4-6 CD and *n* = 5 WD. Unpaired 2-tailed Student’s *t* test was used to test significance. ^#^*P* < 0.1, \**P* < 0.05, \*\*\*\**P* < 0.0001.

We next tested whether maternal exposure to WD influenced fetal liver health. Liver lipids were histologically evident in mWD-exposed fetal livers (**Fig. 2C, upper panel**) and verified by increased LipidSpot staining (**Fig. 2C, lower panel**) and by triglyceride analysis (**Fig. 2D**). Picrosirus red staining, an indicator for fibrosis, was strikingly more evident in mWD fetal liver (**Fig. 2C, middle panel**). Expression of mRNA for genes involved in inflammation (*IL6*), fibrosis (*COL3A1*), and monocyte infiltration (*CCR2*) was elevated in fetal livers from WD-fed dams (**Fig. 2D**), although only *CCR2* expression was significant between groups. Together, these data suggest that exposure to mWD in utero promotes triglyceride storage, a trend for increased inflammation, and early fibrogenesis, which are hallmarks of NAFLD/NASH changes in the fetal liver.

### Maternal WD induces subtle microbiota changes in dams

At 0.6 G, maternal fecal SCFA levels did not differ between diets for propionate and butyrate, but there was an increase in acetate load in the feces of WD-fed dams (**Fig. 3A**). Using 16S sequencing, we found that the maternal gut microbiome was mainly comprised of *Lactobacillales* and *Clostridiales*, with modest changes in microbial composition between groups (**Fig. 3B**). Exposure to WD resulted in significantly lower alpha diversity (**Fig. 4A**) but no community-wide differences between CD and WD groups as measured by beta diversity (PERMANOVA *P* value = 0.26; **Fig. 4B**). Comparing microbiota abundances between diets, feces from mWD dams were enriched in *Acidaminococcus* and unclassified *Betaproteobacteria*, while those from mCD dams were characterized by higher *Anaeroplasmatales* abundance (**Fig. 4C**). Using PICRUSt to predict functional pathways, we found 11 pathways potentially differentiating the effect of diet on the maternal gut microbes. WD-exposed microbiota were predicted to have enrichment of biosynthesis pathways such as gluconeogenesis and aspartate/asparagine synthesis, whereas CD feeding was associated with enrichment of pathways for rhamnose biosynthesis, lactose/galactose degradation, and bacterial-specific biosynthesis of peptidoglycans (*Staphylococci*) and O-antigen (*Escherichia coli*; **Fig. 4C**).

**Figure 3.**
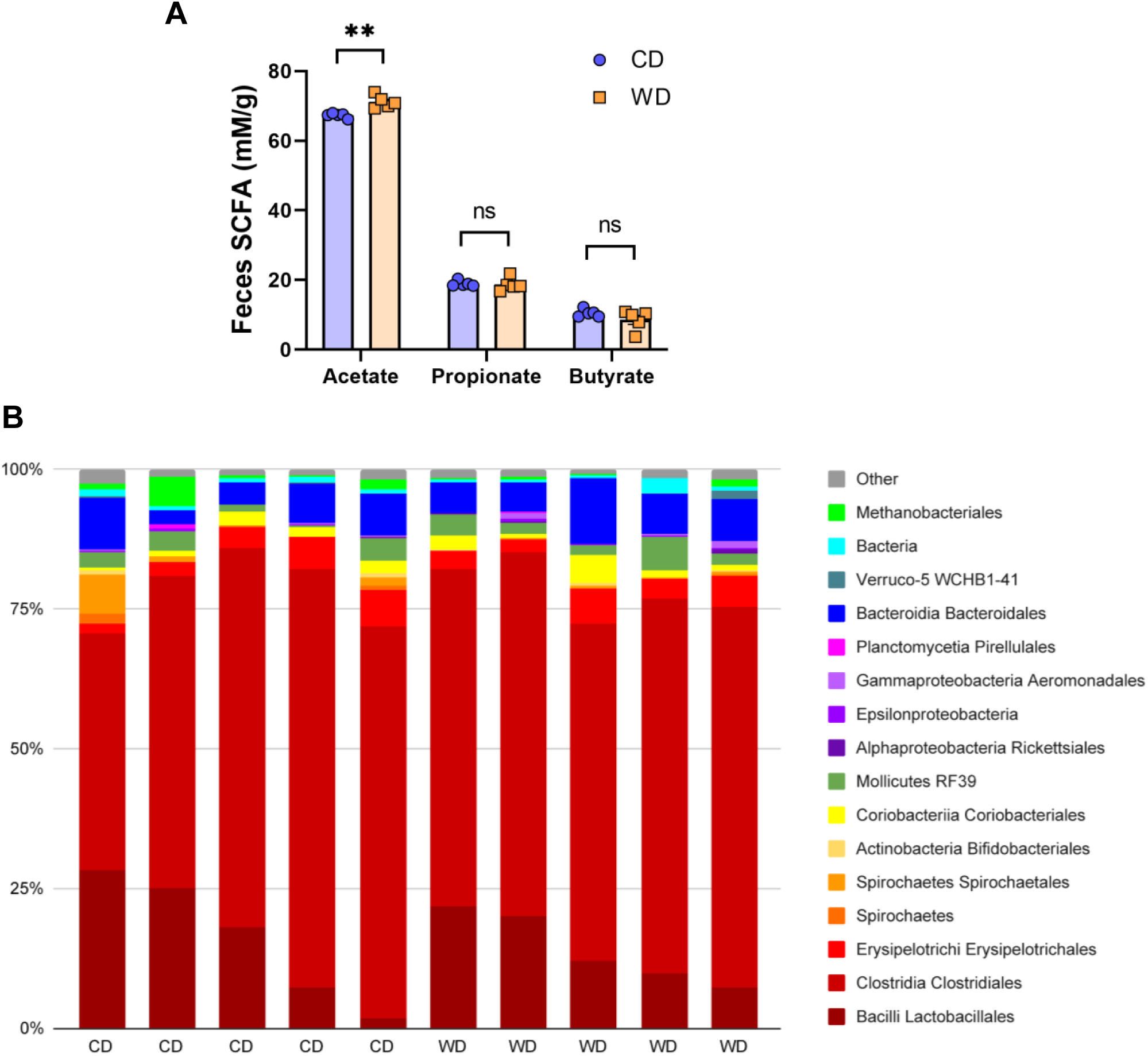
Short-duration exposure to WD induces few alterations in maternal SCFAs and microbiota. (**A**) Abundance of fecal SCFAs. *n* = 5/group. Unpaired 2-tailed Student’s *t* test was used to test significance. \*\**P* < 0.01. (**B**) Microbial abundances for each gut sample, clustered at the order level. Orders comprising less than 0.5% total abundance are displayed as “Other”.

**Figure 4.**
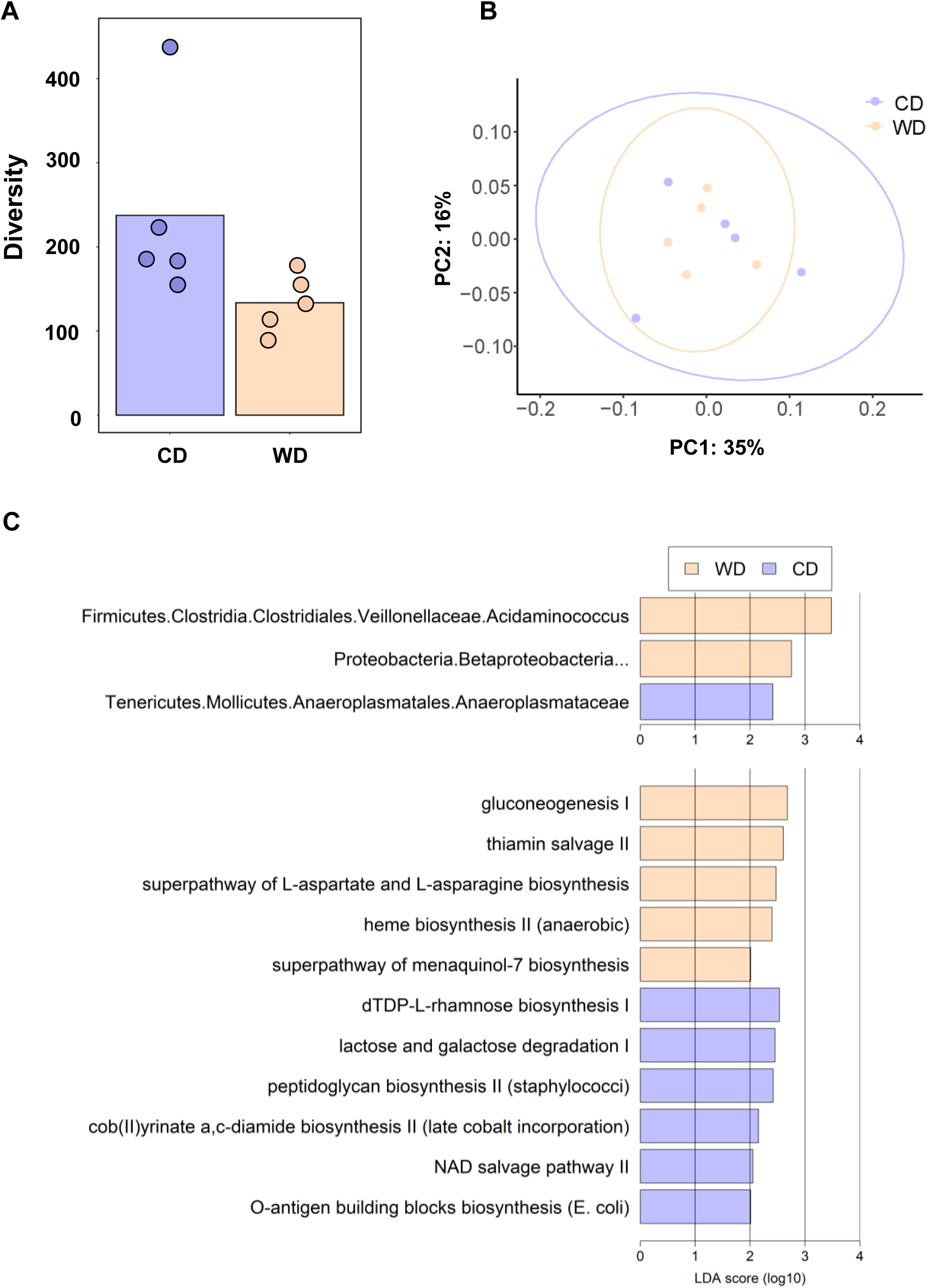
(**A**) Alpha diversity measured using Faith’s phylogenetic diversity. Significance of species richness was tested using the Kruskal-Wallis test (*P* = 0.02). (**B**) PCoA ordination displaying weighted Unifrac beta diversity. Percent variation explained is shown on each axis (PC1: 35% & PC2: 16%). PERMANOVA significance for the weighted Unifrac distances (*P* = 0.26). (**C**) Lefse histograms plotted for significant enrichment in taxa abundances (upper plot) and biochemical pathways (lower plot). *n* = 5/group.

### Maternal WD alters placental and fetal miRNA profiles

Mounting evidence indicates that miRNAs mediate gut microbiome-host molecular communications (70). We sought to explore differences in a targeted set of miRNAs in placenta and fetus associated with maternal diet. We selected miRNAs that were previously found to be differentially expressed in fetal liver in a study in baboons in which the dams were obese, as well miRNAs shown to be regulated by the gut microbiome (miR-122-5p, miR-204-5p, and miR-34a-5p) (71-74) or associated with NASH in rodents and humans (miR-21-5p and miR-29a-3p) (75-78). MiRNA expression analysis in fetal liver tissue (**Fig. 5A**) revealed a significant downregulation of miR-204-5p and miR-145-3p expression upon mWD exposure. In the placenta, we observed a significant downregulation of miR-183-5p and miR-182-5p in the WD-fed group (**Fig. 5B**). In placental tissue, miR-199a-5p showed a trend for downregulation (*P* = 0.057). The expression levels of miR-183-5p, miR-182-5p and miR-199a-5p were directionally similar in both fetal liver and in the placenta, as was the increase in expression of miR-1285-3p.

**Figure 5.**
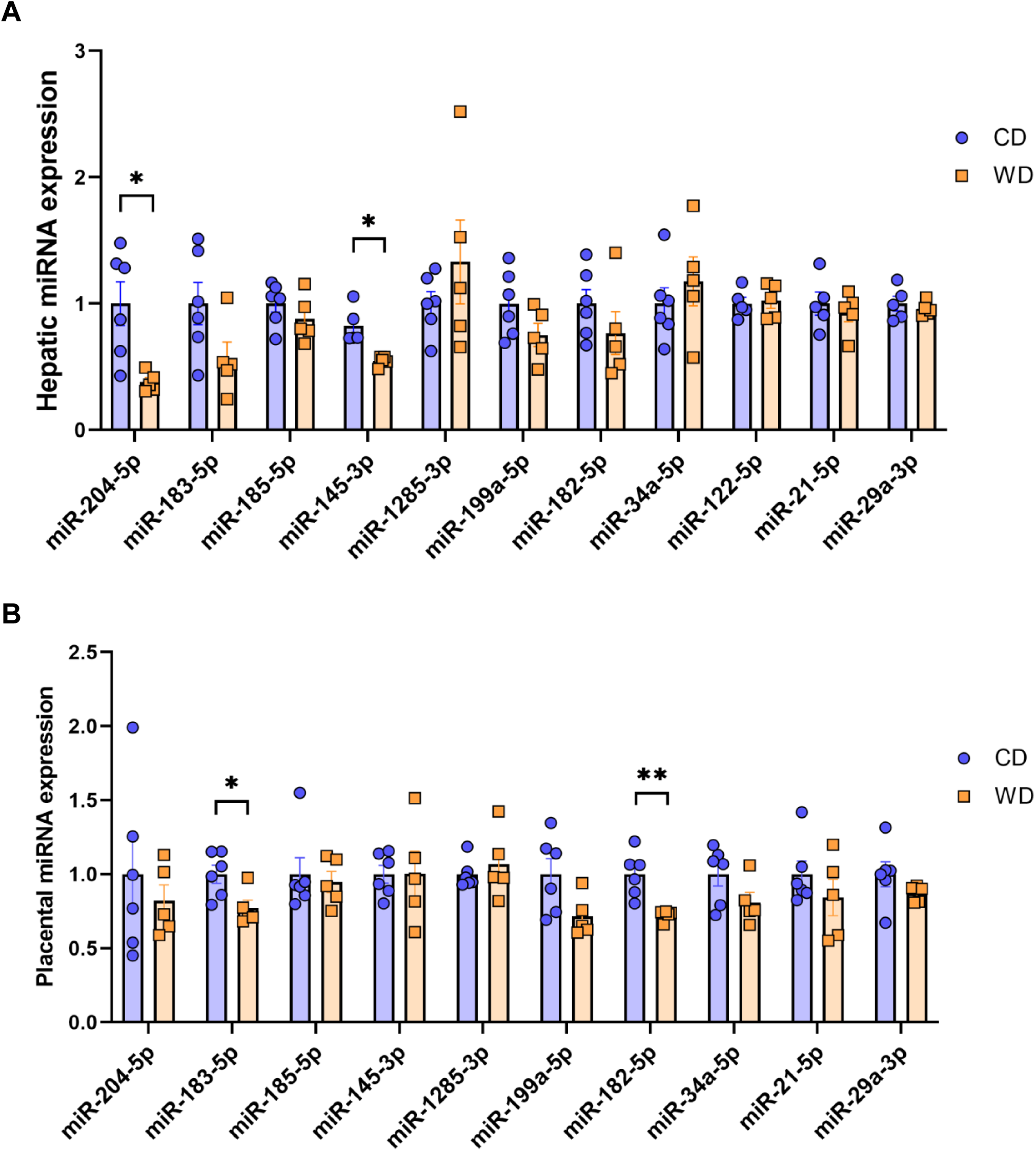
MicroRNA expression analysis. Expression of miRNAs in CD- and WD-exposed fetal liver (**A**) and placental tissue (**B**). *n* = 6 CD and *n* = 5 WD. Unpaired 2-tailed Student’s *t* test was used to test significance. \**P* < 0.05, \*\**P* < 0.01.

### Associations between microbiota, miRNA, metabolic features and fetal liver gene expression

We first analyzed associations between maternal parameters of systemic and lipid metabolism to maternal microbiota and placental miRNAs (**Table 1**). After FDR correction, only unclassified *Rickettsiales* and unclassified *Verrucomicrobia* were significantly and positively associated with maternal HDL and *Desulfovibrionaceae* was positively associated with placental weight. Next, we analyzed associations between maternal parameters of systemic and lipid metabolism and fetal liver TG, miRNA, and mRNA levels (**Table 2**). Fetal liver TGs were positively associated with maternal TGs, HDL, and LDL/VLDL. After FDR correction, only the comparison between fetal liver TGs and maternal LDL/VLDL remained significant. We next compared fetal liver TGs to fetal liver/placental miRNAs and fetal liver mRNA levels (**Table 3**); however, after FDR correction, none of these comparisons were significant. In our analysis of fetal liver expression levels of miRNA and mRNA, we found several placental and fetal miRNAs associated with markers of lipid metabolism, oxidative stress, inflammation, and fibrosis (**Table 4**). The most common miRNA associated with liver mRNA levels was miR-1285-3p. Liver miR-1285-3p was negatively associated with *NFE2L2, ICAM1*, and *VCAM1*, but positively associated with *IL6*. Placental miR-1285-3p was positively associated with *NFE2L2* and negatively associated with *FAP*. After FDR correction, only the association between miR-1285-3p and *IL6* mRNA was significant, though several other associations were trending towards significance (**Table 4**). We next compared fetal liver miRNA levels and relative abundance of maternal microbiota at the taxonomic family level and found miR-122-5p was positively associated with *Succinivibrionaceae* (**Table 5**). Placental miR-1285-3p was positively associated with *Coriobacteriaceae* and *Prevotellaceae*. Both taxa displayed sex differences in association with placental miR-1285-3p, where males remained positively associated (*Coriobacteriaceae*, r2 = 0.57; *Prevotellaceae*, r2 = 0.70) but females showed no association (r2 = 0 for both taxa). After FDR correction, only placental miR-1285-3p remained significantly associated with *Coriobacteriaceae, Prevotellaceae*, and their interactions with sex. Fetal liver miR-122-5p and *Succinivibrionaceae* showed a trend toward association after FDR correction (*P* < 0.1). At the genus level, fetal liver miR-122-5p remained negatively associated with *Blautia* levels and positively associated with *Ruminococcus* after FDR correction. No other genera were selected for inclusion in our LASSO models for any of the other miRNA measured.

**Table 1.** Associations of maternal metabolic measurements compared to maternal microbiota and placental miRNA

|  | Microbiota |  |  |  | Placenta |  |  |  |
| --- | --- | --- | --- | --- | --- | --- | --- | --- |
|  | Taxa (Genus or Family) | r2 | P value | FDR | miRNA | r2 | P value | FDR |
| Dam metabolism |  |  |  |  |  |  |  |  |
| Body wt | *Clostridium* | +0.01 | 0.0374 | 0.5478 |  |  |  |  |
|  | *Oscillospira* | +0.32 | 0.0498 | 0.5729 |  |  |  |  |
|  | Unclassified *Lachnospiraceae* | +0.41 | 0.0040 | 0.1300 |  |  |  |  |
| SSF |  |  |  |  | miR-21-5p | -0.34 | 0.0340 | 0.1831 |
| Dam lipid metabolism |  |  |  |  |  |  |  |  |
| TGs | Unclassified *Bacteroidales* | +0.32 | 0.0029 | 0.1168 | miR-183-5p | -0.50 | 0.0129 | 0.1405 |
|  |  |  |  |  | miR-182-5p | -0.57 | 0.0068 | 0.1368 |
| HDL | Unclassified *Rickettsiales* | +0.43 | <0.0001 | 0.0037 | miR-204-5p:sex | M: -0.56; F: 0.27 | 0.0355 | 0.1834 |
|  | Unclassified *Verrucomicrobia* | +0.38 | 0.0014 | 0.0342 | miR-21-5p:sex | M: -0.70; F: -0.34 | 0.0215 | 0.1565 |
|  | Unclassified *Bacteroidales* | +0.18 | 0.0021 | 0.1156 |  |  |  |  |
|  | *Catenibacterium* | +0.29 | 0.0161 | 0.3247 |  |  |  |  |
|  | *Succinivibrio* | +0.25 | 0.0070 | 0.1890 |  |  |  |  |
| LDL/VLDL | Unclassified *Ruminococcaceae* | +0.22 | 0.0496 | 0.5729 | miR-183-5p | -0.55 | 0.0084 | 0.1368 |
|  | Unclassified *Rickettsiales* | +0.05 | 0.0242 | 0.4329 | miR-199a-5p:sex | M: -0.77; F: 0 | 0.0437 | 0.2007 |
|  | *Treponema* | -0.07 | 0.0122 | 0.2813 |  |  |  |  |
| Dam inflammation |  |  |  |  |  |  |  |  |
| CRP | *Paraprevotellaceae* | +0.49 | 0.0035 | 0.0632 |  |  |  |  |
| Placenta |  |  |  |  |  |  |  |  |
| Weight | *Desulfovibrionaceae* | +0.32 | 0.0012 | 0.0342 |  |  |  |  |
|  | *Methanobrevibacter* | +0.11 | 0.0413 | 0.5547 |  |  |  |  |

Maternal measures were compared to maternal gut microbiota abundances using negative binomial models. Maternal measures were compared to placental miRNAs with ANOVA. Both comparisons utilized LASSO regularization for variable selection. CRP, C-reactive protein; HDL, high-density lipoprotein; LDL, low-density lipoprotein; SSF, sum of skin folds; TG, triglyceride; VLDL, very low-density lipoprotein; wt, weight.

**Table 2.** Associations between maternal and placental characteristics and fetal liver measurements.

|  | Fetal liver | r2 | P value | FDR |
| --- | --- | --- | --- | --- |
| **Dam metabolism** |  |  |  |  |
| Body wt | miR-1285-3p | +0.43 | 0.0171 | 0.1467 |
|  | *VEGFA* | +0.43 | 0.0322 | 0.1816 |
|  | *NFE2L2* | -0.46 | 0.0136 | 0.1405 |
| SSF | *CYP1A1* | -0.57 | 0.0074 | 0.1368 |
| **Dam lipid metabolism** |  |  |  |  |
| TGs | TGs | +0.60 | 0.0088 | 0.1368 |
|  | *ICAM1* | -0.47 | 0.0254 | 0.1577 |
|  | *NFE2L2* | -0.39 | 0.0321 | 0.1816 |
|  | *ACACA* | -0.4 | 0.0403 | 0.1924 |
| HDL | TGs | +0.45 | 0.0207 | 0.1565 |
|  | miR-145-3p | -0.62 | 0.0042 | 0.1368 |
| LDL/VLDL | TGs | +0.87 | 0.0001 | 0.0197 |
|  | miR204-5p | -0.47 | 0.0177 | 0.1467 |
|  | miR204-5p:sex | M: -0.69; F: -0.12 | 0.0464 | 0.2054 |
| **Placenta** |  |  |  |  |
| Weight | miR-34a-5p | +0.64 | 0.0061 | 0.1368 |
|  | *TREM2* | +0.53 | 0.0245 | 0.1577 |
|  | *TLR4* | +0.53 | 0.0254 | 0.1577 |

Maternal measures were compared to fetal liver miRNA and mRNA expression using ANOVA with LASSO regularization for variable selection. HDL, high-density lipoprotein; LDL, low-density lipoprotein; SSF, sum of skin folds; TG, triglyceride; VLDL, very low-density lipoprotein; wt, weight.

**Table 3.** Fetal liver triglyceride associations with miRNAs and mRNAs.

|  | **r2** | **P value** | **FDR** |
| --- | --- | --- | --- |
| Fetal liver miRNA |  |  |  |
| miR-204-5p | +0.38 | 0.0341 | 0.1627 |
| miR-145-3p | +0.54 | 0.0140 | 0.1627 |
| Fetal liver mRNA |  |  |  |
| *COL1A1* | -0.68 | 0.0139 | 0.5819 |
| *CCR2* | +0.51 | 0.0434 | 0.6427 |
| *IL6*:sex | M: -0.91, F: 0.78 | 0.0233 | 0.5819 |
| Placental miRNA |  |  |  |
| miR-182-5p | +0.47 | 0.0168 | 0.1627 |
| miR-34a-5p | +0.39 | 0.0314 | 0.1627 |
| miR-199a-5p:sex | M: -0.94, F: 0.29 | 0.0097 | 0.1627 |
| miR-199a-5p:sex | M: -0.83, F: 0 | 0.0277 | 0.1627 |

Fetal liver triglycerides were compared to fetal liver miRNA/mRNA expression and placental miRNA expression with ANOVA models using LASSO regularization for variable selection.

**Table 4.** Associations between miRNAs and fetal liver mRNA expression levels.

| Pathway/mRNA | Fetal liver<br>miRNA | Placental<br>miRNA | r2 | P value | FDR |
| --- | --- | --- | --- | --- | --- |
| Lipid metabolism/synthesis |  |  |  |  |  |
| *ACACA* | - | miR-183-5p | +0.59 | 0.0096 | 0.0564 |
| *ACACA* | - | miR-182-5p | +0.44 | 0.0305 | 0.0983 |
| *FASN* | - | miR-185-5p | +0.39 | 0.0414 | 0.1132 |
| *FASN* | - | miR-145-3p | -0.54 | 0.0151 | 0.0624 |
| *HMOX1* | - | miR-183-5p | +0.70 | 0.0036 | 0.0520 |
| *SREBP1* | - | miR-185-5p | +0.40 | 0.0394 | 0.1132 |
| Oxidative stress |  |  |  |  |  |
| *NFE2L2* | **miR-1285-3p** | - | -0.50 | 0.0091 | 0.0564 |
| *NFE2L2* | - | **miR-1285-3p** | +0.48 | 0.0107 | 0.0564 |
| Inflammation |  |  |  |  |  |
| *ICAM1* | **miR-1285-3p** | - | -0.63 | 0.0063 | **0.0564** |
| *IL10* | - | miR-183-5p | +0.45 | 0.0288 | 0.0983 |
| *IL10* | **miR-29a-3p** | - | -0.58 | 0.0104 | **0.0564** |
| *IL10* | miR-34a-5p | - | -0.39 | 0.0420 | 0.1132 |
| *IL6* | **miR-1285-3p** | - | +0.76 | 0.0006 | **0.0334** |
| *NCF4* | - | miR-182-5p | +0.48 | 0.0228 | 0.0827 |
| *VCAM1* | **miR-1285-3p** | - | -0.63 | 0.0063 | 0.0564 |
| *VEGFA* | - | miR-185-5p | +0.59 | 0.0093 | 0.0564 |
| *VEGFA* | - | miR-29a-3p | -0.37 | 0.0483 | 0.1167 |
| Stellate cells activation and fibrosis |  |  |  |  |  |
| *COL1A1* | miR-204-5p | - | +0.61 | 0.0132 | 0.0624 |
| *TGFB1* | - | miR-183-5p | +0.69 | 0.0036 | 0.0520 |
| *TGFB1* | - | miR-182-5p | +0.54 | 0.0149 | 0.0624 |
| *FAP* | - | miR-1285-3p | -0.49 | 0.0211 | 0.0814 |
| *TNFSF12* | mir-183-5p | - | +0.38 | 0.0461 | 0.1163 |

Fetal liver mRNA expression was compared to placental and liver miRNA expression using ANOVA models with LASSO regularization for variable selection.

**Table 5.** Associations between maternal microbiota, fetal liver and placental miRNA, and fetal liver mRNA expression levels.

| Microbiota | Liver miRNA | Placental miRNA | Liver mRNA | r2 | P value | FDR |
| --- | --- | --- | --- | --- | --- | --- |
| Family |  |  |  |  |  |  |
| *Succinivibrionaceae* | miR-122-5p |  |  | +0.41 | 0.0384 | 0.0922 |
| *Coriobacteriaceae* |  | miR-1285-3p |  | +0.53 | 0.0105 | 0.0387 |
| *Coriobacteriaceae* |  | miR-1285-3p:sex |  | M: +0.57, F: 0 | 0.0129 | 0.0387 |
| *Prevotellaceae* |  | miR-1285-3p |  | +0.55 | 0.0088 | 0.0387 |
| *Prevotellaceae* |  | miR-1285-3p:sex |  | M: +0.70, F: 0 | 0.0083 | 0.0387 |
| *Clostridiaceae* |  |  | *IL1B* | -0.39 | 0.0326 | 0.0718 |
| *Lachnospiraceae* |  |  | *IL1B* | +0.47 | 0.0175 | 0.0641 |
| *Lachnospiraceae* |  |  | *ACTA2* | +0.40 | 0.0297 | 0.0718 |
| Genus |  |  |  |  |  |  |
| *Blautia* | miR-122-5p |  |  | -0.77 | 0.0011 | 0.0032 |
| *Ruminococcus* | miR-122-5p |  |  | +0.49 | 0.0214 | 0.0321 |

Maternal gut microbiota abundances were compared to fetal liver miRNA/mRNA expression and placental miRNA expression using negative binomial regression models with LASSO regularization for variable selection.

## Discussion

Epigenetic patterning in the placenta and fetus resulting from a Western-style maternal diet may be influenced by maternal gut microbes to promote the development of offspring NAFLD; however, these relationships are poorly described in human-relevant models. Our study using the Olive baboon, in which dams were fed either a CD or WD prior to and during pregnancy, is the first to investigate early associations between the microbiota, placental-fetal miRNAs, and maternal-fetal metabolic dysregulation. Despite ∼1 year of mWD, the dams did not display significant weight gain or adipose deposition, consistent with similar studies in baboons (69) and Japanese macaques (79) where obesity was attained following 2 to 3 years of WD feeding. Strikingly, in the absence of obesity, we found that mWD adversely affected lipid metabolism and inflammation along the maternal-placental-fetal axis, concomitant with decreased miR-182-5p and miR-183-5p in mWD placentas compared with mCD placentas. Previous studies in NHP have not focused on miRNAs in placenta and their association with fetal liver function. We found the reduced placental expression levels of the miR-183 family (including mR-182-5p and miR-183-5p) to be positively associated with expression of a number of NAFLD/NASH-relevant genes in the fetal liver; this family was shown to attenuate pathophysiology in a mouse model of NASH (80) and may therefore be a novel target for future mechanistic studies in our rodent models. We also found significant associations between the maternal microbiota and placental miR-1285-3p, an miRNA associated with promotion of postnatal growth in infants born to mothers with obesity (81). Livers from fetuses of WD-fed dams showed increased steatosis and elevated expression levels of genes involved in macrophage infiltration, inflammation, and fibrosis. These fetal NAFLD indices were increased concomitant with downregulation of hepatic expression of miR-204-5p, an miRNA known to be regulated by gut microbes (74).

Although mechanisms are not fully elucidated, the gut microbiome influences the development of chronic inflammatory diseases such as obesity through a variety of host pathways, including by mediating the expression of host miRNAs (82). The effects of maternal obesity on the offspring microbiome were previously described in Japanese macaques (31, 83), but potential effects of the maternal microbiome on altering epigenetic marks in the placenta or fetus are undetermined. In our model of short-duration WD-feeding, the maternal microbiota were dominated by *Firmicutes*, primarily *Lactobacilliales, Clostridiales*, and *Bacteroidales*, strongly implicated in the maintenance of overall gut function and health (84, 85). In the absence of maternal obesity, we found several associations between the maternal gut microbiota and maternal serum TGs, HDL, LDL/VLDL, and CRP. Similar to our findings associating *Desulfovibrionaceae* abundance with placental weight in baboons, the genus *Desulfovibrio* is enriched in women with GDM (86, 87); notably, GDM is associated with increased placental weight (88). We found placental miR-1285-3p was associated with maternal gut microbiota abundances of *Coriobacteriaceae* and *Prevotellaceae*, both of which displayed a positive relationship with postnatal growth-promoting (81) miR-1285-3p expression in placenta from male fetuses, but no association in females. Both of these bacterial families are metabolically active members of the gut microbiome. MiR-122 accounts for around 70% of all miRNAs in the adult liver and plays a role in regulation of innate immunity (89), proliferation and differentiation of hepatocytes (90), lipid accumulation, and cholesterol metabolism (91). We found a differential association between fetal liver miR-122-5p and two maternal gut genera, *Blautia* and *Ruminococcus*, where *Blautia* negatively associated with miR-122-5p and *Ruminococcus* positively associated with miR-122-5p. Decreased *Blautia* abundance was associated with obesity and intestinal inflammation in children (92) and *Blautia* species abundances have been associated with lowered visceral fat accumulation in human adults (93). Conversely, *Ruminococcus* was positively associated with visceral fat accumulation (94). Determining whether miR-1285-3p and miR-122-5p are early biomarkers for programmed obesity and NAFLD that may be regulated by maternal genera such as *Blautia* and *Ruminococcus* is an important avenue for future exploration.

Compared with mCD dams, mWD dams exhibited dyslipidaemia, characterized by elevated LDL cholesterol and TGs. However, based on the IVGTT at 0.6 G, these dams did not show indices of insulin resistance. These findings are supported by the observations of Short et al., wherein young (5-6 years of age) male baboons fed a WD (high in monosaccharides and saturated fatty acids) for eight weeks exhibited elevated levels of HDL and LDL/VLDL cholesterol and TGs, without a change in body weight or blood glucose (95). Short et al. also noted significant effects on inflammatory indices, with enhanced CD14+ mononuclear cell chemotaxis as well as a significant increase in blood neutrophils (95). We similarly found that mWD dams had mild, systemic inflammation, exemplified by elevated levels of CRP and increased neutrophil counts, which was consistent with some of the changes in cytokine profiles known to be exacerbated by maternal obesity. These observations suggest that upregulation of at least some pro-inflammatory programs during pregnancy are mediated by the diet alone.

Our finding of a trend for increased numbers of macrophages in the placentas of WD-fed dams is suggestive of enhanced chemoattraction of maternal monocytes or fetal monocytes/Hofbauer cells to the placenta, or activation of maternal peripheral monocytes that target the placenta, and is consistent with previous findings of monocyte priming and enhanced monocyte chemotaxis in male baboons fed a WD for eight weeks (95). Peripheral monocytes from obese pregnancies displayed elevated chemokine receptor expression and enhanced migration capacity (23). Further, placental resident CD14+ and CD68+ mononuclear cells (macrophages) increased 2-to 3-fold in obese pregnancies (23, 96). Previously, Frias and colleagues noted increased placental inflammatory cytokine expression and placental infarctions in the Japanese macaque model of chronic WD feeding and maternal obesity (97). We did not specifically address monocyte priming in the current study; however, we did note that mRNA expression of both *TNF* and *IL8* mRNA was elevated mWD placentas, consistent with WD-induced ‘priming’ of placental inflammation (in combination with recruitment of macrophages). We also did not find notable gross placental pathologies (calcifications, infarcts) in mWD placentas compared with mCD placentas.

Our novel study has several strengths including our use of the Olive baboon as a translational model for developmental programming. An advantage over rodents, baboons are similar to humans in gestational duration, placentation (hemochorial monodiscoid), singleton births, hormone profiles, and social behaviors. Further, our model allowed us to address the role of mWD independent from maternal obesity in nulliparous females. One drawback of this study is the small sample size which limited the conclusions we could draw from our analyses. In a baboon model where maternal obesity was attained, Puppala et al. showed a trend toward increased hepatic lipid accumulation and more severe steatosis, assessed histologically, that did not progress toward NASH in the fetus (48). This is in contrast to our findings and those of Wesolowski et al. (12) and McCurdy et al. (79) in obese Japanese macaques where fetuses from WD-fed dams had higher hepatic TGs associated with impaired mitochondrial function and increased fibrogenesis, which Nash et al. demonstrated was localized to the hepatic periportal region (98).

In contrast to our findings using a targeted analysis, Puppala et al. used an untargeted microarray-based approach and found that miR-145-3p was upregulated in livers from baboon fetuses exposed to maternal obesity and associated with a decrease in SMAD4; they additionally reported an increase in miR-182-5p and -183-5p (48). Our findings are also consistent with observations previously reported in obese Japanese macaques where a differential response to high-fat diet in dams was found, allowing segregation into insulin-sensitive and insulin-resistant subgroups (99). Maternal insulin resistance (elevated TGs, insulin, and weight gain) led to activation of de novo lipogenic and pro-inflammatory pathways in offspring liver at 1 year of age (99). Moreover, Elsakr et al. showed that prolonged WD feeding, multiple diet switches, and increasing age and parity were associated with increased insulin resistance in dams (100). The dams in our study, subjected to a relatively short-duration exposure to WD, were not insulin resistant and did not have significant weight gain compared with matched CD-fed dams; although the effects we observed in the fetal liver were modest, they are striking given the maternal exposures were limited.

We conclude that, prior to the onset of obesity, a WD initiated several months preceding gestation and maintained over its course causes perturbed maternal lipid homeostasis and impacted maternal gut microbiota composition. Both maternal metabolic parameters and maternal gut microbiota were associated with expression of fetal liver miRNA and mRNA that are markers for lipid metabolism, oxidative stress, and inflammation, suggesting that maternal diet, in the absence of obesity, has significant consequences for epigenetic regulation of fetal and infant health.

## Conflict of Interest

The authors declare that the research was conducted in the absence of any commercial or financial relationships that could be construed as a potential conflict of interest.

## Author Contributions

JP, JF, KJ, and DM contributed to conception and design of the study. KS, AM, RJ, MC-C, and KJ contributed to writing the original draft of the manuscript. SG, JP, and DM collected maternal data. RJ, SG, MD, and RB performed experiments. KS, MT, and DD performed microbiome and association analyses. DD, M-PA, JF, and DM contributed to supervision of the study. All authors contributed to the article and approved of the submitted version.

## Funding

Funding for this study was received from the Harold Hamm Foundation/Presbyterian Hospital Foundation, grant 20211471 (DM, JF, KJ, JP, DD, M-PA).

## Acknowledgments

We thank the OUHSC Stephenson Cancer Tissue Pathology Core, supported partly by NIGMS P20GM103639 and NCI P30CA225520, for the liver histology. We thank for OUHSC Laboratory for Molecular Biology and Cytometry Research core for library construction and sequencing.

## Data Availability Statement

Data supporting this study will be made available by the corresponding author, KJ, upon request. 16S sequencing data is available from the NIH Sequence Read Archive (Accession number in progress).

## Notes

### Competing Interest Statement

The authors have declared no competing interest.

