## Supplemental Table 1 for "Western diet-induced shifts in the maternal microbiome are associated with altered microRNA expression in baboon placenta and fetal liver"

Supplementary Material

Western diet-induced maternal microbial dysbiosis is associated with steatosis and altered expression of microRNAs in baboon fetal liver and placenta

Kameron Y. Sugino^1,†^, Ashok Mandala^1,†^, Rachel C. Janssen^1^, Sunam Gurung^2^, MaJoi Trammell^3^, Michael W. Day^3^, Richard S. Brush^4^, James F. Papin^5^, David W. Dyer^3^, Martin-Paul Agbaga^4,6^, Jacob E. Friedman^1,7^, Marisol Castillo-Castrejon^7,8^, Karen R. Jonscher^1,9,*^ and Dean A. Myers^2^

| **Supplemental Table S1.** qPCR primer sequences for mRNA expression. | | |
| --- | --- | --- |
| **Gene** | **Forward Primer** | **Reverse Primer** |
| *ACACA* | AGCCCTCAACAAAGTCCTCG | AGTGGGTCACCCCATTGTTG |
| *ACTA2* | GGCAAGTGATCACCATCGGA | GTGGTTTCATGGATGCCTGC |
| *ACTB* | GGGAAATCGTGCGTGACATT | AGGTAGTTTCGTGGATGCCA |
| *CCR2* | TTGACGTGAAGCAAATCGGG | CCAGCATGTTGCCCACAAAA |
| *COL1A1* | AAGGACAAGAGGCACGTCTG | CAGGAAGGTCAGCTGGATGG |
| *COL3A1* | GGTCCAAAGGGTGACAAGGG | GGCCAGGAGGACCAATAGGA |
| *FAP* | AGTTTCAGCGACTACGCCAA | TCTTGATCAGTGCGTCCGTC |
| *FASN* | GAGCACAACAGGGTGCTAGA | TTGATGATCAGGTCCACGGC |
| *HMOX1* | AGCAACAAAGCGCAAGACTC | GATCCATCGGAGAAGCGGAG |
| *ICAM1* | CAGTTCGTGCTGAAAGCCAC | ATCGGGGTCCATACAGGACA |
| *IL10* | CTGAGAACCACGACCCAGAC | GAAGAAATCGATGACAGCGCC |
| *IL1B* | GCTCTCCACCTCCAGGGACAGG | TGAGGCCCAAGGCCACAGGT |
| *IL1B* * | TGAAAGCTCTCCACCTCCAG | TTGGGCAGACTCGAATTCCA |
| *IL6* | CAGTTCCTGCAGAAAAAGGCAA | GAGATGCGTCGTCATGTCCT |
| *IL6* * | ATGCAATAACCACCCCTGAA | CTGCAGCCACTGGTTCTGT |
| *IL8* * | CCTTTCCACCCCAAATTTATC | TTCTGTATTGACGCAGTGTGG |
| *MCP1 (CCL2)* | TTCTGTGCCTGCTGCTCATAG | GGGGGCATTGATTGCATCTGG |
| *NCF4* | ATACCTGCCCTCAACGCCTA | CGGGGACACACTCTTGACTT |
| *NFE2L2 (NRF2)* | TCTGCCAACTACTCCCAGGT | AACGTAGCCGAAGAAACCTCA |
| *RPS9* | TTGAAGCTGATCGGCGAGTAT | TCTCATCAAGCGTCAGCAGTT |
| *SREBF1 (SREBP1)* | GGATTGCACGTTCGAAGACAT | AGAGAGGAGCTCAGTGTGGT |
| *TGFB1* | GAGCCCTGGACACCAACTAC | GAGGTCCTTGCGGAAGTCAA |
| *TLR4* | GCTTCCTCCGTTTTCCAGAACTGC | TGGAGAGGTGGCTTAGGCTCTGA |
| *TNF* * | TTCAGCTGGAGAAGGGTGAT | CCAAAGTAGACCTGCCCAGA |
| *TNFSF12* | CAGGACCCATCGGAACTGAA | CCGTGTTTTCCGGCCTTTAG |
| *TREM2* | ATCTACAACCCCACGATGCG | CAGAGATCTCCAGCATCCCG |
| *VCAM1* | TTTTTGTCAATGTTGCCCCC | CAGGCTGTAGCTCCCCATTAG |
| *VEGFA* | CCCACTGAGGAGTCCAACAT | CTCCTATGTGCTGGCCTTGG |

### * Denotes qPCR primer set used in placental tissue.
